# Structural basis for targeting the human T-cell leukemia virus Tax oncoprotein and syntenin-1 interaction using a small molecule

**DOI:** 10.1101/2021.08.25.457680

**Authors:** Sibusiso B. Maseko, Inge Van Molle, Karim Blibek, Christoph Gorgulla, Julien Olivet, Jeremy Blavier, Charlotte Vandermeulen, Stéphanie Skupiewski, Deeya Saha, Thandokuhle Ntombela, Julianne Lim, Frederique Lembo, Aurelie Beauvois, Malik Hamaidia, Jean-Paul Borg, Pascale Zimmermann, Frank Delvigne, Luc Willems, Johan Van Weyenbergh, Dae-Kyum Kim, Franck Dequiedt, Haribabu Arthanari, Kourosh Salehi-Ashtiani, Steven Ballet, Alexander N. Volkov, Jean-Claude Twizere

## Abstract

Human T-cell leukemia virus type-1 (HTLV-1) is the causative agent of adult T-cell leukemia (ATL). Although ATL is a well-characterized T-cell neoplasm, linked to intermittent expression of the viral Tax-1 protein, there is currently no strategy to target Tax-1 functions using small molecules. Here, we report a comprehensive interaction map between Tax-1 and human PDZ domain-containing proteins (hPDZome). We show that Tax-1 interacts with more than one-third of hPDZome components, including proteins involved in cell cycle, cell-cell junctions, cytoskeleton organization, and membrane complex assembly. Using nuclear magnetic resonance (NMR) spectroscopy, we have determined the structural basis of the interaction between the *C*-terminal PDZ binding motif (PBM) of Tax-1, and the PDZ domains of syntenin-1, an evolutionary conserved hub that controls exosome trafficking. Finally, we have used confocal imaging, molecular modelling, NMR and mammalian cell-based assays to demonstrate that the Tax-1/syntenin-1 interaction is amenable to small-molecule inhibition. Altogether, our study highlights the biological significance of Tax-PDZ interactome and its interplay with exosome formation. It shows a direct link between extracellular vesicles and HTLV-1 transmission, providing a novel framework for the design of targeted therapies for HTLV-1-induced diseases.

## INTRODUCTION

Human T-cell lymphotropic viruses (HTLV-1 to −4) are members of *Deltaretrovirus* genus of the *Retroviridae* family. As the first human retrovirus isolated 40 years ago (Poiesz et al., 1980), HTLV-1 is the etiological agent of adult T-cell leukemia/lymphoma (ATL), an aggressive neoplasm (Hinuma et al., 1981), as well as HTLV-1-associated myelopathy (HAM), also referred to as tropical spastic paraparesis (TSP), a fatal neurological disease (Gessain et al., 1985). Currently, about 10-20 million people worldwide are infected with HTLV-1, and about 5% will develop severe ATL or HAM/TSP diseases (Legrand et al., 2022; Pasquier et al., 2018; Rowan et al., 2020). HTLV infections are endemic in Southern Japan, central Africa, the Caribbean region, South America, and in central Australia (Aboriginal communities)(Einsiedel et al., 2021; Legrand et al., 2022; Willems et al., 2017). For the last 40 years, ATL patients have been treated by means of chemotherapy-based approaches, but with very limited benefit, as the median survival rate (8-10 months) does not change through chemotherapy (Bazarbachi et al., 2011; Cook et al., 2019; Cook and Phillips, 2021; Nasr et al., 2017). Improved survival is, however, achieved by antiviral therapies combining zidovudine and interferon-alpha, allogenic hematopoietic stem cell transplantation, or the use of monoclonal antibodies targeting the CC chemokine receptor 4 (CCR4), which is frequently expressed in ATL patient samples (Cook and Phillips, 2021; Fuji et al., 2016; Ishida et al., 2015; Yoshie et al., 2002). However, the above therapies are not very effective, mainly due to clinical disease heterogeneity, treatment discrepancies between countries, and a lack of specific and universally targeted drugs (Hleihel et al., 2021). In contrast to ATL, no treatment is available for HAM/TSP patients.

In addition to the essential retroviral genes *gag, pol*, and *env*, HTLV-1 encodes regulatory proteins, including Tax-1, Rex, p12I, p13II, p30II, and HBZ. Among these, Tax-1 and HBZ are the main viral factors contributing to the disease. For example, both proteins were shown to induce leukemia in transgenic mouse models independently (Hasegawa et al., 2006; Satou et al., 2011). Additionally, the expression of both proteins may induce an inflammatory process in the spinal cord during the development of TSP/HAM (Casseb and Penalva-de-Oliveira, 2000). However, there is a difference in kinetics between the expression of Tax-1, which is heterogeneous and intermittent, and the expression of HBZ, which is more stable (Kulkarni and Bangham, 2018; Miura et al., 2019). These proteins also perturb the global quantitative and qualitative transcriptome of host cells in an opposing manner (Vandermeulen et al., 2021). Regarding the Tax-1 protein, it is a potent transcriptional activator of viral and cellular genes through association with transcription modulators, which are directly involved in T-cell proliferation, migration, apoptosis, and cell-cycle (reviewed in Boxus et al., 2008; Enose-Akahata et al., 2017; Giam, 2021; Martinez et al., 2019; Ratner, 2020).

Tax-1 is a multidomain protein harboring intrinsically disordered linker regions, encompassing the segments 76-121, 252-275, and the *C*-terminal amino acid stretch 320-353 (Boxus et al., 2008). This modular organization ensures conformational plasticity and potentially explains the dynamic hijacking of crucial cellular functions through interaction with a multitude of interacting partners (Simonis et al., 2012; Vandermeulen et al., 2021; Wu et al., 2004). The carboxyl terminus of Tax-1 contains a PDZ-binding motif (PBM), which confers binding to a class of proteins embedding a defined structure of ~90 amino acids known as PDZ domains (PSD-95/Discs Large/ZO-1) (Hirata et al., 2004; Makokha et al., 2013; Takachi et al., 2015; Tsubata et al., 2005; Xie et al., 2006). PDZ-containing proteins are modular and implicated in the assembly of large protein complexes, mediating signaling and cell polarity and communication (Harris and Lim, 2001; Stevens and He, 2022). PDZ domains contain a globular fold characterized by five to six antiparallel β-strands and two α-helices in a βA-βB-βC-αA-βD-βE-αB-βF arrangement (Christensen et al., 2019). Even though there is structural information about PDZ domains, there are still conflicting issues regarding their binding selectivity and thus their classification (Tonikian et al., 2008). Early studies have classified them into three distinct classes (classes I, II, and III) based on the affinity to their binding partners. Class I recognizes a PBM with T/S-X-Ψ whilst classes II and III recognize Ψ-X-Ψ and D/E-X-Ψ respectively, where X is any amino acid, and Ψ, a hydrophobic amino acid (Christensen et al., 2019; Tonikian et al., 2008). A recent analysis of the human genome estimates the presence of 271 PDZ domains in 154 distinct proteins (human PDZome), excluding variants and isoforms (Amacher et al., 2020). To date, 14 PDZ proteins interacting with HTLV-1 Tax have been identified (Boxus et al., 2008; Pérès et al., 2018; Simonis et al., 2012). These Tax-1-PDZ interactions have important implications in the HTLV-1-induced leukemogenesis process. In particular, it was shown that the Tax-1 PBM motif, which is not present in the HTLV-2 Tax counterpart, promotes the transformation of rat fibroblasts, mouse lymphocytes *in vitro* (Higuchi et al., 2007; Hirata et al., 2004; Tsubata et al., 2005), as well as the persistence of the HTLV-1 virus, and proliferation of T-cells *in vivo* (Pérès et al., 2018; Xie et al., 2006).

In this study, we hypothesized that targeting Tax-1-PDZ interactions would provide a gateway towards novel anti-HTLV-1 transmission strategies as new means to combat ATL. Therefore, we comprehensively mapped the Tax-1 interactome with human PDZ-containing proteins and found that about one-third of the human PDZome is capable of specific interactions with Tax-1. Using nuclear magnetic resonance (NMR) spectroscopy, we structurally characterized interactions between Tax-1 PBM and PDZ domains of syntenin-1, a “seed protein” in small extracellular vesicle formation and trafficking (Leblanc et al., 2020; Lim et al., 2021). Finally, we demonstrate that FJ9, a small molecule able to disrupt the Tax-1-syntenin-1 interaction, impedes Tax distribution in the Golgi apparatus and virological synapses and inhibits several Tax-1 functions, including cell immortalization and HTLV-1 cell-to-cell transmission.

## MATERIAL AND METHODS

### Predicting potential Tax interactors using *in silico* motif-domain and domain-domain based strategies

We combined two approaches, similar to Zanzoni et al. (Zanzoni et al., 2017), to predict potential human host targets of Tax-1. These were: i) Motif-domain based interaction inferences, if Tax-1 possesses a short linear motif (SLiM), m, that binds to a complementary domain d in protein A. For instance, with Tax having a PBM, if a given protein contains a PDZ domain, it is inferred as one of the potential interactors of Tax, and ii) domain-domain based interaction inferences in which Tax and a human host interactor protein A are predicted to interact with each other if Tax possesses an interaction domain d1 (e.g., the Zinc finger-like domain in Tax-1) that binds with interaction domain d2 (e.g., another zinc finger-like domain) in host interactor protein A. Hence, a protein with zinc finger-like domain would be inferred as one of the potential interactors of Tax. All the templates for inferring motif-domain and domain-domain based interactions were retrieved from the following resources a) the 3did Database (Mosca et al., 2014) b) ELMdb (Dinkel et al., 2012) c) Garamszegi et al. (Garamszegi et al., 2013). As such, 535, 232, and 666 motif-domain template interactions were retrieved from 3did, ELMDb, and Garamszegi et al., respectively. Finally, an exhaustive list of 1311 motif-domain based template interactions was prepared combining the three sources. Domain-domain and motif-domain interaction templates were downloaded from the 3did database, which currently stores 6290 high-resolution interaction interfaces.

### Identification of Short linear motifs (SLiMs) in the Tax protein sequence

Short linear motifs in the Tax sequence were searched using SLiMSearch 2.0 tool from SLiMSuite (Davey et al., 2011). SLiMSearch 2.0 takes as input a set of motif sequences and tests them against queried protein sequences. The motif consensus is represented as a regular expression and contains ordinary and special characters. Permitted ordinary characters are single-letter amino acid codes. The special characters are based on regular expression syntax as defined in the Eukaryotic Linear Motif resource (Kumar et al., 2020). To reduce the number of false positives, only those motifs which fall in disordered regions of the Tax sequence were retained. In order to detect motifs situated in the disordered regions of the Tax sequence, each motif was assigned a disorder score (DS_c_):

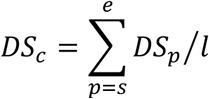

Where, DS_p_ is the IUPred disorder score of the residue at position p in the Tax protein sequence, s is the start position of the consensus match, e is the end position of the consensus match, l is the consensus match length. Motif instances whose disorder score was greater than 0.4 were retained (Zanzoni et al., 2017). The rationale behind selecting motif instances located in the disordered region is that the functional SliMs are largely located in the unstructured region of the protein (Edwards et al., 2012; Fuxreiter et al., 2007).

### Cell culture

The HEK and HeLa cell lines were cultured in DMEM (Dulbecco’s Modified Eagle’s Medium) containing 10% FBS (Fetal Bovine Serum), 10% glutamine, and antibiotics (Penicillin 100U/ml, Streptomycin 100U/ml). The lymphocyte cell lines (JURKAT, JURKAT-LTR, JPX9, CEM, CTV1, MOLT16, HUT78) were maintained in culture in RPMI medium (Roswell Park Memorial Institute medium), 10% glutamine, and antibiotics (Penicillin 100U/ml, Streptomycin 100 μg/ml). The lymphocyte cell lines infected with HTLV-1 (MT2, MT4, HUT102, C8145, C91PL) were cultured under the same conditions as the lymphocyte cell lines not infected with HTLV-1, in addition to the conditions of containment (biosafety) level L3. All cells were incubated at 37°C with 5% CO_2_.

### Transient transfections

HEK cells cultured to 80% confluency in DMEM were transfected using polyethylenimine (PEI). The medium was changed before transfection, and cells were collected 24 hours after transfection. HeLa cells were transfected using the Lipofectamine 2000 reagent (Invitrogen) according to the manufacturer’s instructions and collected 24 hours after transfection. Cells transfected with siRNAs were cultured in DMEM culture medium at 40-50% confluence. siRNA transfection was performed with calcium phosphate using the ProFection® Mammalian Transfection System (Promega) transfection kit according to the manufacturer’s instructions. The culture medium was changed 24 hours post transfection and cells were harvested 48 hours post transfection.

### Tax-1 Yeast 2 Hybrid with individual PDZ Domains

The Gateway cloning method was used to transfer DNA encoding the PDZ domain into the AD yeast expression vector pACT2. All PDZ domains were transformed into the yeast haploid strain Y187 (MATa, ura3-52, his3-200, ade2-101, leu2-3, 112, gal4Δ, met-, gal80Δ, MEL1, URA3: GAL1UAS -GAL1TATA -lacZ). Similarly, the DNA fragment encoding Tax-1 was cloned into the DB yeast expression vector pGBT9 and transformed next into the haploid yeast strain AH109 (MATa, trp1-901, leu2-3, 112, Ura3 −52, his3-200, gal4Δ, gal80Δ, LYS2: GAL1UAS-GAL1TATA-HIS3, GAL2UAS-GAL2TATA-ADE2, URA3: MEL1UAS-MEL1TATA-lacZ, MEL1). The interactions between each PDZ and Tax-1 were tested by conjugation of the two strains of yeast. Briefly, overnight culture of the two types of yeast (of the opposite mating type) was performed. This in a selective medium favoring diploid yeast (liquid yeast extract-Peptone-Dextrose YPAD –W Tryptophan –L Leucine) supplemented with 10% PEG for 4 h at 30 ° C and *via* gentle stirring. After washing with water, the yeasts were placed on a solid Tryptophan-Histidine-Leucine (-WHL) selective medium for phenotypic assay. Diploid yeasts which proliferate on the -WHL medium indicate an interaction between the two fusion proteins.

### GST-pulldown

To detect Tax-1 partners by pulldown, PDZ domains were expressed in HEK293 cells. GST-Tax-1, or GST-Tax-2 protein as negative control, were expressed separately and then purified with glutathione Sepharose beads. Beads containing the fusion proteins were then incubated with PDZ-flag containing cell lysate. The detection of Flag by immunoblot was indicative of an interaction between Tax-1 and the PDZ domain protein.

### Gaussia princeps luciferase complementation assay (GPCA)

HEK293 cells were seeded in 24-well plates and transfected next with GL1 and / or GL2 plasmids expressing the fusion proteins 24 hours later. Luciferase activity was measured on the lysates in 96-well plates. The results were normalized relative to the value of the Luciferase control Renilla. The normalized luciferase ratio was calculated as follows: NLR = co-transfection luciferase value (GL1 + GL2) / (GL1 luciferase value alone + GL2 luciferase value alone). An interaction was considered positive or validated when NLR ≥ 3.5. Cell lysis and luminescence measurements were performed in triplicate for each condition.

### Measurement of Luciferase activity

Collected cells were lysed with the passive lysis buffer (Promega). For luciferase measurements, 100 μl of the lysates were used, to which the substrates Firefly and Renilla stop & glo (Promega) were added. Luciferase measurements were performed in 96-well plates using an automated DLR machine. Firefly luciferase values were normalized to Renilla luciferase values, and the calculated ratio represents luciferase activity.

### Synthetic Peptide

The Tax-1 10-mer (H-SEKHFRETEV-OH) peptide was synthesized by Biomatik, Canada, as the acetate salt of >98% purity.

### Expression and purification

Plasmids harboring the PDZ domains of syntenin-1 (PDZ1, PDZ2 and tandem PDZ1+PDZ2) were purchased from GeneScript and transformed into *E. coli* BL21. A single bacterial colony was used to inoculate 5.0 mL LB and grown overnight at 37 °C, from which 1.0 mL was used as a starter culture for expressing the proteins in 100 mL LB. For expression, the culture was grown at 37°C, shaking (180rpm) until OD600 reached 0.4-0.6, then induced by the addition of IPTG to a final concentration of 1.0 mM and further grown for 4 hrs under the same conditions. The isotopically labeled U-[^13^C,^15^N] or U-[^15^N] proteins were produced in *E coli* grown in the minimal medium following a published protocol (Volkov et al., 2013). GST-fused proteins were purified using glutathione 9epharose 4B beads (GE HealthCare) equilibrated in 1x Phosphate Buffered Saline (PBS) buffer. Bound proteins were eluted with 50 mM Tris, pH 8, 10 mM reduced glutathione. GST tag was then cleaved off using preScission protease (Sigma) overnight at 4°C. The mixture was then loaded back to the column equilibrated in PBS, and PDZ proteins were collected from the flow-through. These were further purified using a HiPrep 16/60 Sephacryl S-100 HR size exclusion column, equilibrated in 20 mM NaPi pH 6.0, 50 mM NaCl, 2 mM DTT.

### Nuclear Magnetic Resonance Spectroscopy

All NMR spectra were acquired at 298 K in 20 mM sodium phosphate 50 mM NaCl pH 6.0, 2 mM DTT and 10 % D_2_O for the lock on a Bruker Avance III HD 800 MHz spectrometer (equipped with a TCI cryoprobe) or a Varian Direct-Drive 600 MHz spectrometer (fitted with a PFG-Z cold probe). The data were processed in NMRPipe (Delaglio et al., 1995) and analyzed in CCPNMR (Vranken et al., 2005). The assignments of backbone resonances of the individual syntenin-1 PDZ1 and PDZ2 domains were obtained from 0.7-1.1 mM U-[^13^C,^15^N] protein samples using a standard set of 3D BEST HNCACB, HN(CO)CACB, HNCO and HN(CA)CO experiments and aided by the published assignments (Cierpicki et al., 2005; Liu et al., 2007; Tully et al., 2012). Subsequently, the NMR assignments of the individual PDZ domains were transferred to the spectrum of syntenin-1 PDZ1+2 tandem and verified by standard triple-resonance experiments.

The chemical shift perturbation experiments and NMR titrations were performed by incremental addition of 10 mM stock solutions of Tax peptide to U-[^15^N] or U-[^13^C,^15^N] protein samples at the initial concentration of 0.3 mM. At each increment, changes in chemical shifts of the protein resonances were monitored in 2D [^1^H,^15^N] HSQC spectra. The average amide chemical shift perturbations (Δδ_avg_) were calculated as Δδ_avg_ = (Δδ_N_^2^/50+Δδ_H_^2^/2)^0.5^, where Δδ_N_ and Δδ_H_ are the chemical shift perturbations of the amide nitrogen and proton, respectively. For the **FJ9** binding experiments, 4 μl of 50 mM **FJ9** stock solution in d_6_-DMSO (or pure d_6_-DMSO for the reference sample) was added directly to 200 μl of 0.1 mM U-[^15^N] protein solution, and [^1^H,^15^N] HSQC spectra were acquired in 3 mm NMR tubes with the total sample volume of 204 μl,

The NMR titration curves were analyzed with a two-parameter nonlinear least squares fit using a one-site binding model corrected for the dilution effect (Kannt et al., 1996), Equation 1 :

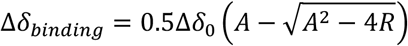

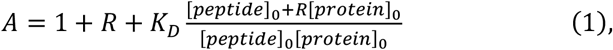

where Δδ_binding_ is the chemical shift perturbation at a given protein/peptide ratio; Δδ_0_ is the chemical shift perturbation at 100 % peptide bound; R is the [peptide]/[protein] ratio at a given point; [protein]_0_ and [peptide]_0_ are the concentrations of the starting protein sample and the peptide titrant stock solution, respectively; and K_D_ is the equilibrium dissociation constant.

### RNA interference

The double-stranded interfering RNAs (siRNAs) were purchased from Eurogentec (Liège, Belgium). The following siRNAs were used in this work:SDCBP sense (5 ‘-GCAAGACCUUCCAGUAUAA-3’), SDCBP anti-sense (5’ - UUAUACUGGAAGGUCUUGC-3’), and control siRNA (5’ - GGCUGCUUCUAUGAUUAUGtt-3’).

### Immunoblots

The cells were lysed and clarified by centrifugation. After incubating the lysate for 5 min at 100°C in the presence of Laemmli buffer, electrophoresis was carried out on 10% SDS-PAGE, followed by immunoblotting. The following antibodies were used: anti-Flag, anti-SDCBP, anti-tax and anti-ubiquitin.

### Immunofluorescence and confocal microscopy

HeLa cells were seeded on coverslips in 24-well plates for 48 hours. The cells were then washed three times at 37 °C with PBS, fixed with PBS-4% paraformaldehyde for 15 min, and then washed twice more with PBS. Cells were then permeabilized with PBS-0.5% Triton X-100 for 20 min, incubated with the PBS-FBS 20% blocking solution for 30 min, and then washed twice with PBS. Subsequently, the cells were incubated with the corresponding primary antibody diluted in PBS-Triton X-100 0.5% for 2 hours, washed three times with PBS, and incubated 1 hour with the corresponding Alexa conjugated secondary antibodies (Invitrogen), diluted 1/1000 in PBS-Triton. Finally, the cells were washed three times with PBS and mounted on glass coverslips with ProLonf Gold Antifade containing DAPI (Life technologies). The slides were examined by confocal microscopy with a Leica TCS SP2 or Nikon A1R confocal microscope, equipped with a 60X objective. The images were taken with a resolution of 1024×1024 pixels and subsequently processed and assembled with LEICA LAS AF Lite 5 or A1R Software.

### qRT-PCR

Total RNA was extracted using the GeneJET RNA purification kit (Thermo scientific). Total RNA was reverse transcribed with random primers using the RevertAid First Strand cDNA Synthesis Kit (Thermo Scientific). QPCR was performed using Roche’s SYBER Green detection in the Lightcycler 480 (Roche). Quantification of mRNA was performed by relative normalization to the constitutive gene GAPDH. The relative expression levels were calculated for each gene using the ΔCt or ΔΔCt method.

### Purification of exosomes

Exosomes were isolated from cell supernatants. First, 50-100 ml of the supernatants were centrifuged at 2000 xg for 20 minutes at room temperature to remove floating cells. Second, the cell-free supernatant was then centrifuged at 12,000 xg for 45 minutes and filtered (0.22 μm) to remove additional cell debris. Third, the samples were ultracentrifuged for 2 h at 100,000 g 4°C. The pellet containing the exosomes was washed with PBS, then centrifuged for an additional 2 hours at the same speed, and finally suspended in lysis buffer (50 mM Tris-HCl, 1% SDS, pH 7.5, supplemented with protease and phosphatase inhibitors). The lysate was stored at 4°C until use in Western blot.

### Tax-1 / SDCBP immunoprecipitation

The interaction of Tax-1 with SDCBP was re-validated by immunoprecipitation. HEK 293T cells were transfected with Flag-Syntenin, Tax-1 and Tax-2. Forty-eight hours after the transfection, the cells were lysed and protein expression verified by a Western blot, using specific anti-Tax-1 and anti-Flag antibodies.

### Cellular localization of Tax-1 and SDCBP

HeLa cells were transfected with constructs expressing Tax-1 and control siRNA or SDCBP siRNA. Cells were fixed, permeabilized, and labeled with primary anti-Tax-1 antibodies and secondary antibodies conjugated to Alexa 488 24 hours post-transfection. These were then analyzed under a confocal microscope. Using Nikon A1R software, we quantified the intensities of ROI (regions of interest), which allowed to determine the cytoplasm / nucleus ratio of protein localization. Statistically significant data are indicated with * (P <0.05), ** (P <0.01), or *** (P <0.001).

### Mutual effect of overexpression of Tax-1 and SDCBP

HeLa cells were transfected with constructs expressing Tax-1 and / or Flag-SDCBP. The cells were fixed, permeabilized and labeled with anti-Tax-1 or anti-Flag antibodies and conjugated secondary antibodies Alexa 488 or Alexa 633 24 hours post transfection. These were then analyzed under a confocal microscope as described for the previous section.

### Molecular Docking

The crystal structures of the syntenin-1 PDZ1 and PDZ2 domains were obtained from the protein data bank (PDB) entry 1W9E (Grembecka et al., 2006). The ions and water molecules were removed and the proteins prepared by adding hydrogen atoms using the Chimera software (Pettersen et al., 2004). The **FJ9** compound was prepared by assigning bond orders and atomic charges with the Gasteiger method using antechamber (Gasteiger and Marsili, 1980). The substrate center coordinates (X = 29.65, Y = 9.97, and Z = 63.49 Å) in the PDZ binding site were used to generate a cubic grid box (X = 21.98, Y = 18.58, and Z = 16.29 Å), large enough to accommodate the ligand. The AutoDock Vina (Trott and Olson, 2010) embedded in Chimera was used to dock the **FJ9** compound. The resulting solutions were ranked according to their energy scores relative to the root-mean-square deviation (RMSD) with a cut-off of < 2 Å. The highest-ranked docking poses were visually inspected according to a set of guideline criteria (Fischer et al., 2021) and those with the ligand binding poses engaging the canonical PDZ binding site were selected for further analysis. The docking scores for the final PDZ1-**FJ9** and PDZ2-**FJ9** complexes were −7.2 kcal/mol and −6.8 kcal/mol, respectively, with the corresponding RMSD values of 1.19 Å and 1.25 Å.

### Tax-1-induced transformation inhibition by FJ9

Rat-1 cells were stably transduced (rat fibroblasts) with a vector lentiviral pTax-1-IB to test the effect of **FJ9** on the transformation of cells induced by the HTLV-1 virus. A pool of resistant cells (5 × 10^3^) was seeded in DMEM / 10% FCS containing 0.33% agarose, superimposed on a layer of 0.5% agarose in a 6-well plate with 100 μM of **FJ9** or DMSO. After three weeks in culture, colonies that usually form “foci” were examined under a light microscope and photos taken to quantify the effect of **FJ9**. The Tax-1 induced foci count was performed using ImageJ software.

### Test for inhibition of cell-to-cell transmission of Tax-1 by FJ9

Virus-producing cells (MT2) were co-cultured with cells containing the gene encoding luciferase under the control of the HTLV-1 LTR5’ viral promoter. The cells were grown for 24 hours in the presence or absence of 100 μM of **FJ9**. Luciferase activation assay was performed using a firefly luciferase kit (Promega E1500).

### Test for inhibition of the interaction of Tax-1 with SDCBP by FJ9

The fluorescent protein YFP was used to estimate the ability of **FJ9** to inhibit the interaction between Tax-1 and SDCBP. HEK 293T cells were transfected with the *C*-terminal fragment of the YFP protein fused to SDCBP and with the *N*-terminal fragment of YFP fused to Tax-1. The cells were then treated with 300 μM **FJ9** or DMSO 24 hours later and further cultured for an additional 24 hours before flow cytometry analysis. The results represent the mean and standard deviation of three independent experiments.

### Statistical analysis

The values of the graphs are presented as the mean ± standard deviation, calculated on at least three independent experiments. Significance was determined using a t test with two variables (comparison of means). The thresholds of value P (p value) are represented as follows; * = p <0.05; ** = p <0.01 and *** = p <0.001

## RESULTS

### A comprehensive interactome map of Tax-1 with human PDZ-containing proteins

The *Tax-1* viral oncogene is a hallmark of the initiation and maintenance of HTLV-1-induced diseases. However, the encoded Tax-1 protein is still viewed as “undruggable”, essentially because of its pleiotropic effects and low expression levels in most HTLV-1-infected cells. We reasoned that a detailed understanding of the Tax-1 interactome could reveal specific Tax-1 interactome modules, amenable to small molecule inhibition. We have previously highlighted the function of Tax-1 as a *“hub”* molecule, capable of interacting with several host cell complexes at the transcription, post-transcription, and translation levels (Boxus et al., 2008; Simonis et al., 2012; Vandermeulen et al., 2021). To date, both high throughput and small-scale/focused studies have identified 258 human proteins interacting with Tax-1 (Vandermeulen et al., 2021). However, to comprehensively estimate the size of the Tax-1 interactome, we identified potential interacting partners based on motif-domain and domain-domain interactions. As such, we predicted 2401 interactions with human proteins (PPIs), including 168 experimentally validated PPIs with Tax-1 (Figure 1A). This suggests that about 10% of the full human proteome could be affected by this viral oncoprotein through a well-defined set of interaction modules (Table S1).

**Figure 1.**
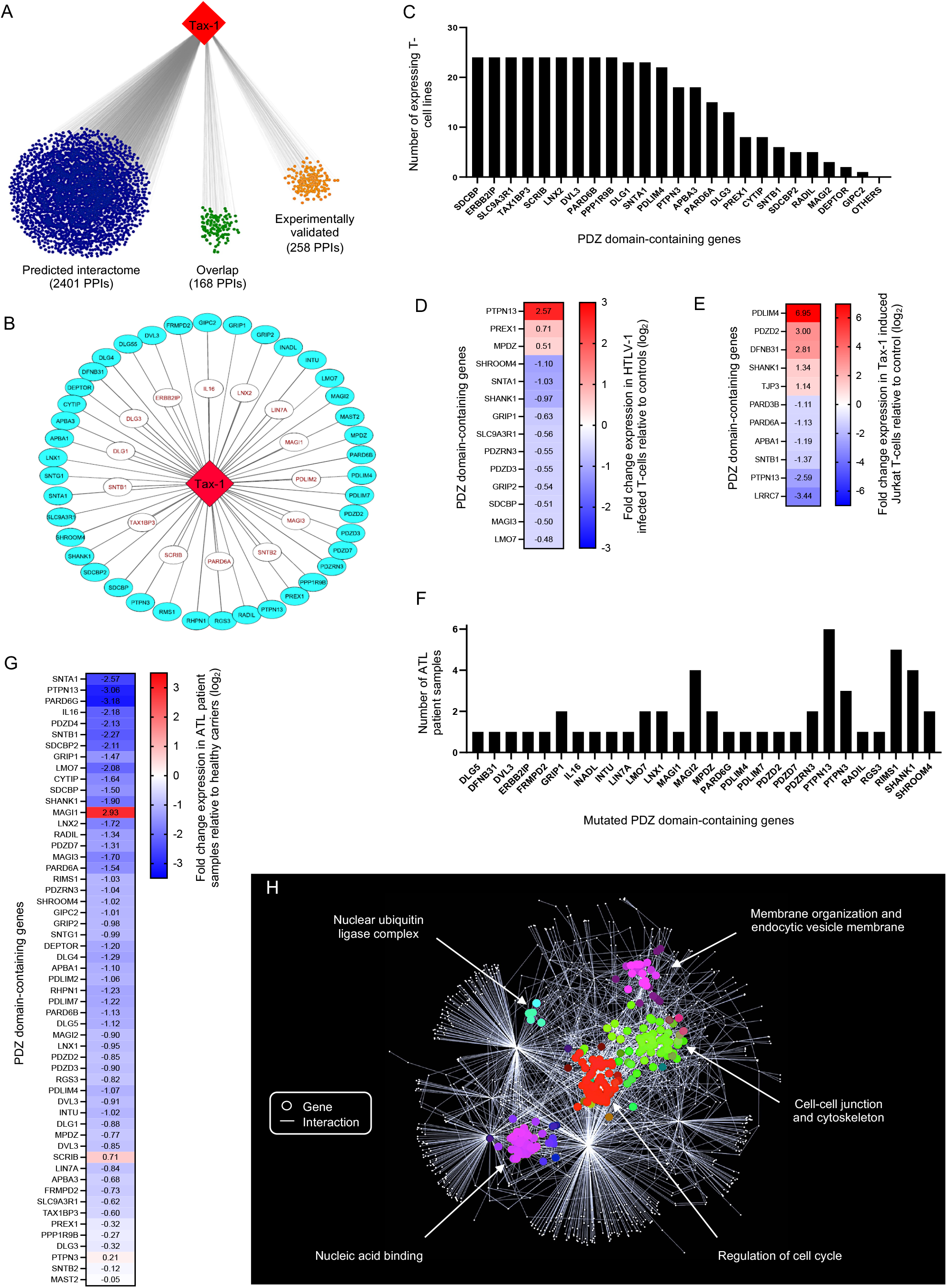
A comprehensive analysis of the Tax-1 interactome with human PDZ-containing proteins. (A) *In silico* prediction of interactions between Tax-1 domains and motifs and human protein domains listed in Table S1. (B) A map of Tax-1-PDZ interactome. Red indicates overlap with known PPIs from the literature. (C) Expression of the interacting PDZ-containing protein genes across 24 different T-cell lines. (D) Differential expression data in HTLV-1-infected versus non-infected T-cells (genes for which p<0.05 are shown). (E) Differentially expressed PDZ genes in Jurkat T-cells following induction of the Tax-1 expression (genes for which p<0.01 are shown). (F) PDZ genes mutated in ATL patient samples. (G) Differentially expressed PDZ genes in ATL patient samples relative to healthy carriers (genes for which p<0.05 are shown). (H) SAFE analysis of the Tax-1/ human PDZ proteins interactome. See also Figure S1, Tables S1

In order to understand the functional plasticity of the Tax interactome, we hypothesized that transient associations controlled by multi-domain scaffolding proteins are able to dynamically relocate Tax-1 complexes within the cell, as demonstrated for binary interactome yeast models (Lambourne et al., 2021). In the context of Tax-1, human PDZ-containing proteins are ideal targets because the vast majority of them are hub proteins with several intracellular partners (Table S2) located in different cellular compartments where Tax-1 exerts its perturbation functions (Giam, 2021; Vandermeulen et al., 2021). To generate a complete interactome of Tax-1 with human PDZ-containing proteins, we combined three different sources of Tax-1 protein-protein interaction (PPI) datasets (Figures 1B and S1A), including (i) literature-curated high quality interactions (Figure S1B), (ii) systematic yeast two-hybrid interactions obtained by testing a library of 248 individualized PDZ domains, orthogonally validated using the Gaussia princeps luciferase complementation (GPCA) assay (Cassonnet et al., 2011); and (iii) GST-pulldown analyses of open reading frames (ORFs) available in the human ORFeome collections (http://horfdb.dfci.harvard.edu/hv7/) (Figures S1C-D).

Our approach allowed us to generate a Tax-PDZ interactome (Figure 1B) encompassing 54 human proteins. This represents 35% of the entire human PDZome (Amacher et al., 2020), and a substantial increase from the 14 already known human PDZ proteins directly interacting with Tax (Figure S1B).

To further determine the relative implication of PDZ-containing proteins in the human interactome, we used our recently released human binary reference interactome map (HuRI) (Luck et al., 2020) and ranked PDZ proteins according to their number of direct interactors (“first degree” or proteins interacting with Tax-1 partners) or indirect associations (“second degree”) (Table S2). The average number of first and second degree Tax-1 partners represented 24 and 536 PPIs, respectively, further emphasizing the scaffolding role of PDZ-containing proteins targeted by the viral Tax-1 oncoprotein.

To assess the overall biological significance of the Tax-PDZ interactome, we compared our new dataset with transcriptome and genomic data of human PDZ protein-coding genes in different relevant datasets: (i) expression data across 24 different T-cell lines (Figure 1C); (ii) differential expression data in HTLV-1-infected versus non-infected cells (Figure 1D); (iii) differential expression data in Jurkat cells expressing Tax-1 versus control cells (Figure 1E); (iv) mutational landscapes in ATL patient samples (Figure 1F); and (v) transcriptome data from a “gene signature” of ATL (Bazarbachi, 2016; Fujikawa et al., 2016) (Figure 1G). Taken together, this comprehensive analysis highlights the implication of PDZ genes and their products in the biology of HTLV-1. Finally, we performed a spatial analysis of functional enrichment (SAFE) (Baryshnikova, 2016) using the above datasets and a STRING network of Tax-1/PDZ interactors (Table S2). We identified five significantly enriched biological processes, including nucleic acid binding, nuclear ubiquitin-binding, regulation of cell cycle, cell-cell junction and cytoskeleton organization, and endocytic vesicle membrane assembly (Figure 1H). The identified modules suggest that, through interaction with PDZ proteins, Tax-1 can affect multiple cellular processes, which may be needed for the viral functions at different disease progression stages.

### Crosstalk between Tax-1-PDZ interaction and membrane vesicles trafficking

The above SAFE analysis highlighted endocytic vesicle membrane assembly as one of the enriched functions in the Tax-1/PDZ interactome (Figure 1H). The subnetwork of this function highlights syndecan binding protein 1 (SDCBP1 also known as syntenin-1), which has the highest connectivity in this subnetwork (Figure 2A) and also in our recently released human interactome map (Luck et al., 2020) (HuRI, http://www.interactome-atlas.org/). Syntenin-1 is particularly relevant for viral infections as it controls extracellular vesicle (EV) formation and cell-to-cell communication(Imjeti et al., 2017; Leblanc et al., 2020; Lim et al., 2021). Importantly, Tax-1, which is a regulator protein, has been shown to localize into EVs from HTLV-1 infected cells. These Tax-1-containing EVs significantly contribute to viral spread and disease progression (Al Sharif et al., 2020; Pinto et al., 2019). However, the molecular mechanisms controlling Tax-1 recruitment into EVs are unknown. We, therefore, tested the possibility that syntenin-1 could mediate Tax-1 localization into exosomes. As shown in Figure 2B, a siRNA targeting syntenin-1 expression is associated with a reduction of Tax-1 levels in EVs, in accordance with Tax-1 translocation from the cytoplasm to the nucleus (Figure 2C). These findings suggest that syntenin-1 is required for the presence of Tax-1 in EVs. We have extensively characterized the Tax-1/syntenin-1 interaction, as well as the interaction between Tax-1 and syntenin-2, using yeast two hybrid (Figure 2D), co-immunoprecipitations (Figure 2E), and GST-pulldown assays (Figure 2F). It appears that both syntenin proteins (SDCBP and SDCBP2) interact with Tax-1, but not with HTLV-2 Tax, lacking a PBM, further demonstrating the specificity of Tax-1’s PBM for syntenin PDZ domains. Thus, thanks to its PDZ domains, syntenin-1 can act as a molecular bridge, targeting Tax-1 to EVs *via* the ESCRT pathway.

**Figure 2.**
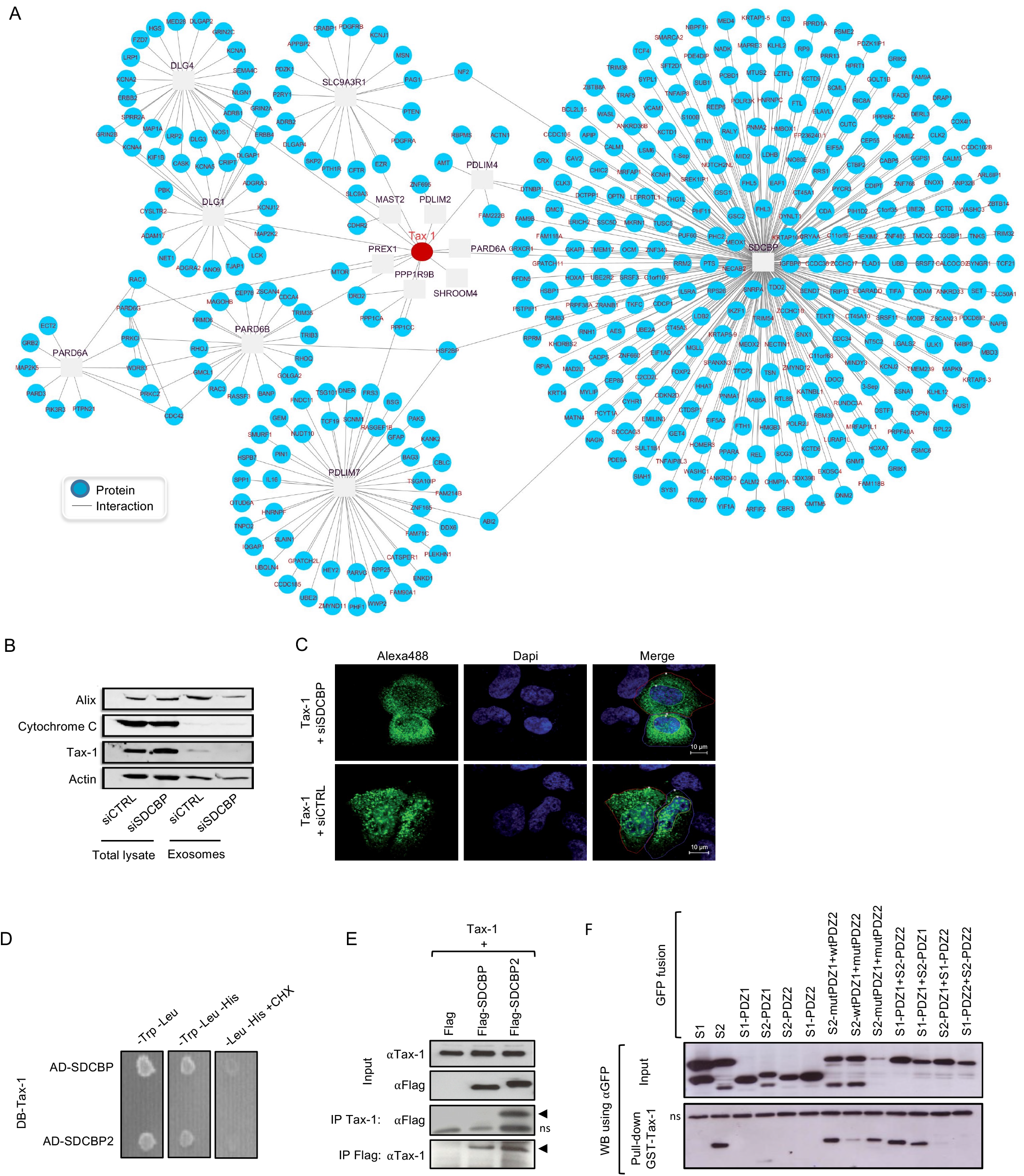
A crosstalk between Tax-1-PDZ interactions and membrane vesicles trafficking. (A) Tax-1 interactome with the cytoskeleton. (B) HEK293 cells transfected with Tax in the presence of a control siRNA or SDCBP siRNA. Exosomes were purified and characterized by western blot using indicated antibodies, in comparison with whole cell lysates. (C) Confocal microscopy examination of HeLa cells expressing Tax-1 in the presence of a control siRNA or SDCBP siRNA HTLV-1 Tax (Tax-1). Bar graphs indicate relative quantification of the Alexa-fluor 488 intensities. (D) Tax-1 fused to the Gal4 DNA-binding (DB) domain, and SDCBP or SDCBP2 fused to Gal4 activating domain (AD) interact in yeast. (E) Tax-1 co-immunoprecipitates with SDCBP and SDCBP2 in transfected HEK293 cells. (F) Pulldown of indicated SDCBP (S1) and SDCBP2 (S2) domains fused to GFP using GST-Tax-1 bound to glutathione beads.

### The structural basis of Tax/ syntenin-1 PDZ interactions

To determine the structural basis of Tax-1/syntenin-1 interaction, we employed solution NMR spectroscopy. Monitored in a series of [^1^H,^15^N] heteronuclear single-quantum correlation (HSQC) experiments, binding of the natural-abundance, NMR-silent *C*-terminal Tax-1 decapeptide (abbreviated as Tax-1 10-mer) to the ^15^N labeled, NMR-visible PDZ1, PDZ2, or tandem PDZ1+2 domains of syntenin-1 causes large spectral changes (Figure 3). Addition of Tax-1 10-mer to ^15^N syntenin-1 PDZ1+2 leads to incremental chemical shift changes for some of the resonances, several of which progressively disappear in the course of the titration, which is symptomatic of the fast-to-intermediate NMR exchange regime. The chemical shift perturbations map to the canonical peptide binding sites of both PDZ1 (including residues I125 and G126) and PDZ2 (residues V209, G210, F211 and F213) domains, encompassing the β1/β2 loop, the β2 strand, and the α2 helix (Figure 3A). The binding effects are stronger for the PDZ2 domain and appear to extend to the back of the protein (including residues T200, L232 and T266) (Figure 3A,B). These findings are consistent with an earlier study of syntenin-1-peptide interactions by X-ray crystallography that highlighted a binding-induced reorientation of the α2 helix (Grembecka et al., 2006), which could explain the chemical shift perturbations of the residues at the back of PDZ2 (Figure 3B).

**Figure 3.**
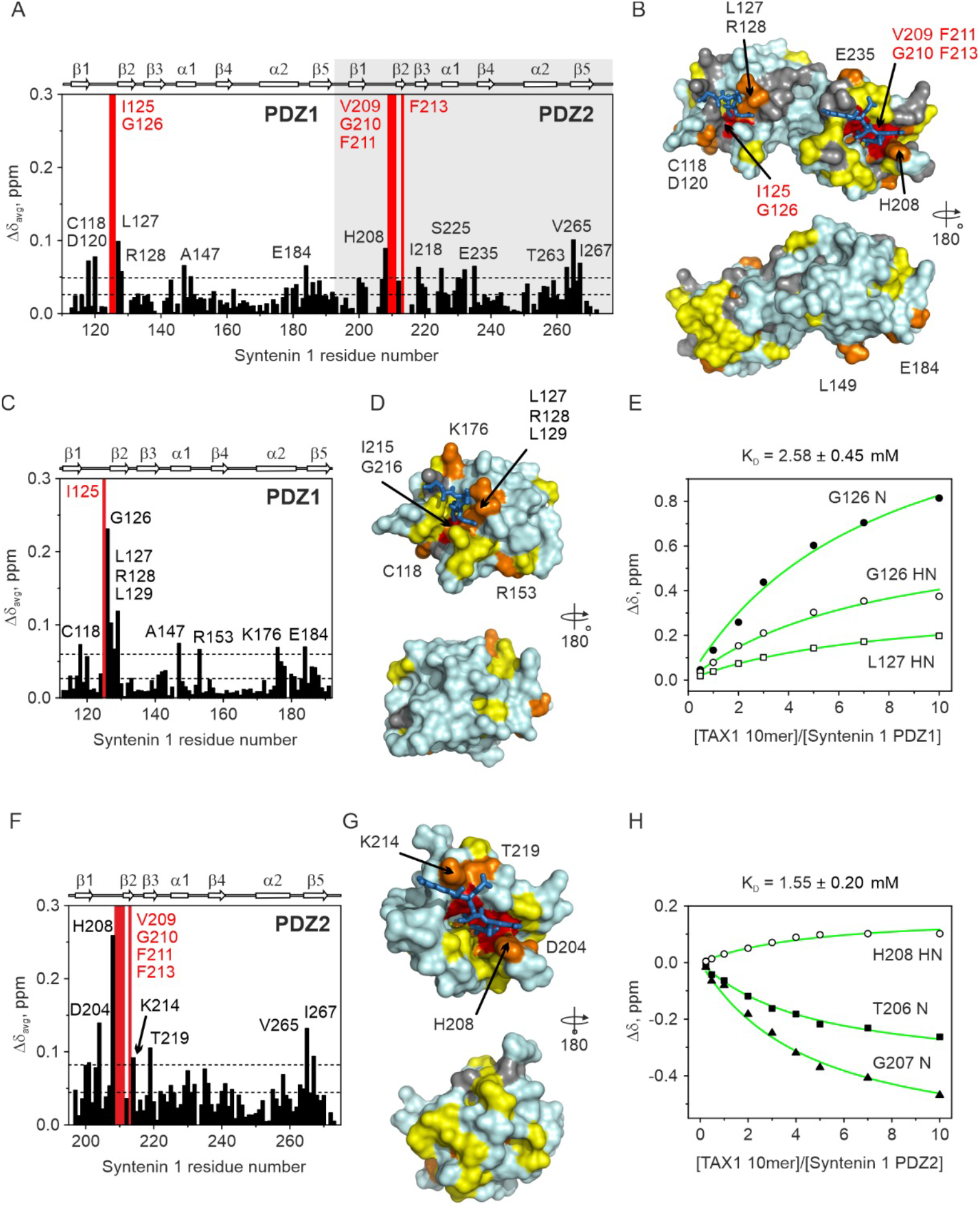
NMR-observed binding of the Tax-1 10mer peptide to syntenin-1 PDZ domains. The interaction of Tax-1 10mer with ^15^N syntenin-1 (A-B) PDZ1+2 tandem or individual (C-E) PDZ1 and (F-H) PDZ2 domains. (A,C,F) Δδ_avg_ in the presence of 5 molar equivalents of Tax-1 10mer. Residues with backbone amides broadened beyond the detection limit upon complex formation are indicated by red bars. The horizontal lines show the average Δδ_avg_ and the avg + stdev. Residues with large Δδ_avg_ are labelled. The secondary structure of syntenin-1 PDZ1 and PDZ2 domains, drawn from the PDB entry 1W9E(Grembecka et al., 2006b), is shown above the plot. (B,D,G) Chemical shift mapping of the Tax-1 10mer binding. The molecular surfaces of syntenin-1 PDZ domains is colored by the Δδ_avg_ values (yellow: Δδ_avg_ > avg; orange: Δδ_avg_ > avg + stdev; red: broadened out upon binding). Prolines and residues with unassigned or strongly overlapping backbone amide resonances are in grey. The modeled C-terminal parts of the Tax-1 peptide, bound to the canonical PDZ sites, are shown in sticks. (E,H) NMR chemical shift titrations with the Tax-1 10mer peptide. Chemical shift perturbations of the backbone amide atoms indicated in the plots were fitted simultaneously to a binding model with the shared K_D_ (Equation 1). The solid lines show the best fits with (E) K_D_ = 2.58 ± 0.45 mM for syntenin-1 PDZ1 and (H) K_D_ = 1.55 ± 0.20 mM for syntenin-1 PDZ2.

To evaluate the binding contribution of each PDZ domain, we performed NMR experiments with the Tax-1 10-mer and individually expressed syntenin-1 PDZ1 and PDZ2 domains. Overall, the chemical shift perturbation maps for isolated PDZ1 (Figure 3C-E) and PDZ2 (Figure 3F-H) are similar to those for the PDZ1+2 tandem (Figure 3A-B). A slight preference for the PDZ2 domain is observed, with K_D_ values of 2.6 and 1.6 mM for PDZ1 and PDZ2, respectively (Figure 3E,H).

### FJ9 is a small molecule binder of syntenin-1 PDZ domains

To determine the extent to which small molecules could bind syntenin-1 PDZ domains and potentially interfere with Tax functions, we tested **FJ9**, a previously characterized small molecule inhibitor of Frizzled-7 Wnt receptor, and the PDZ domain of Dishevelled (Dsh) (Fujii et al., 2007). Addition of **FJ9** to the individual ^15^N-labeled syntenin-1 PDZ1 and PDZ2 domains leads to chemical shift perturbations in 2D [^1^H, ^15^N] HSQC spectra (Figure 4A,E). **FJ9** binding affects syntenin-1 residues in and around the canonical peptide binding sites: in particular I125 and G126 of PDZ1 (Figure 4B,C) and V209 and G210 of PDZ2 (Figure 4F,G). These are the same amino acids that interact with the Tax-1 10-mer peptide (Figure 3), suggesting that **FJ9** can serve as a competitive inhibitor of Tax-1/syntenin-1 interaction.

**Figure 4.**
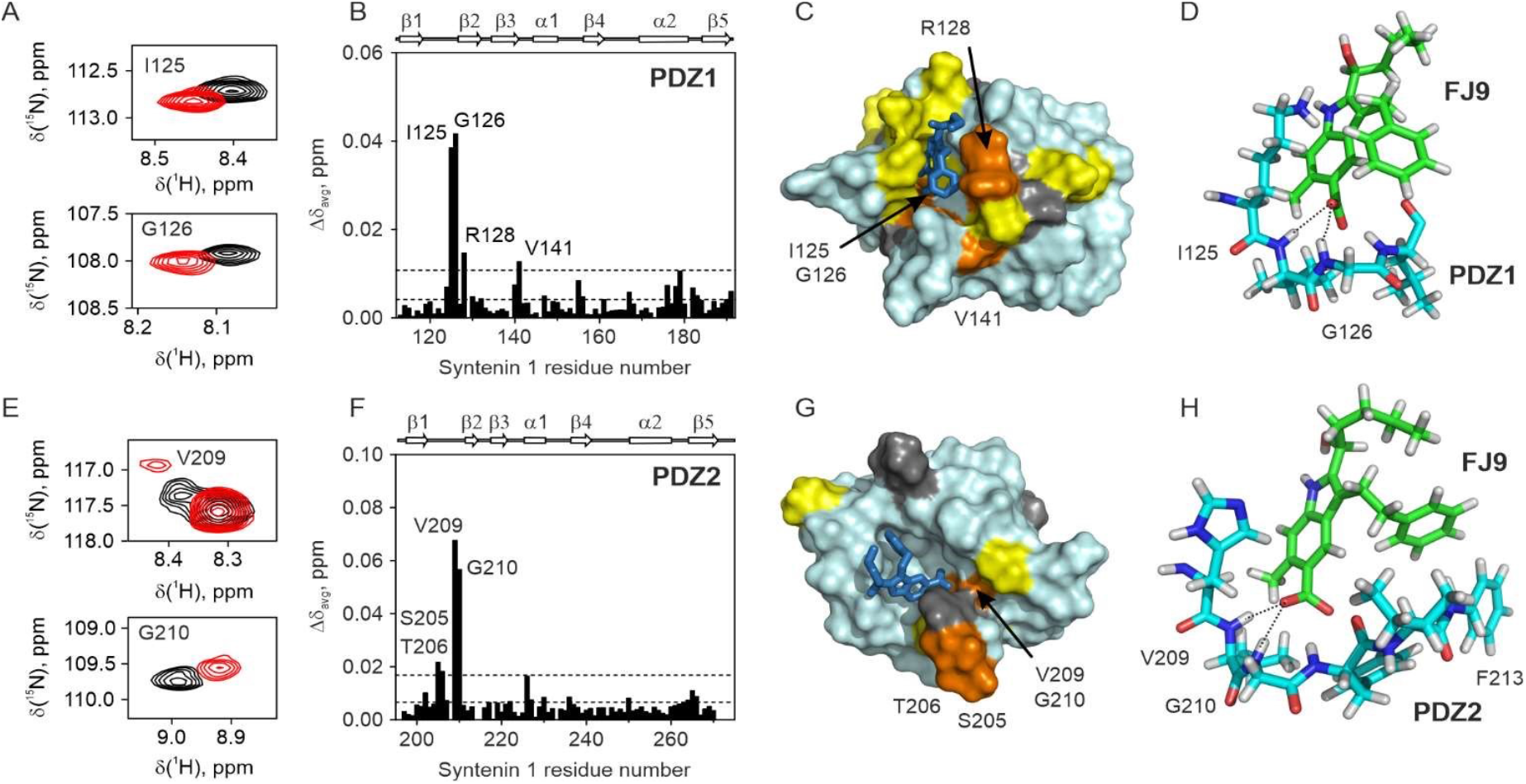
FJ9 binding to syntenin 1 (A-D) PDZ1 and (E-H) PDZ2 domains. (A, E) Selected regions of ^1^H-^15^N HSQC spectra of the free (black) or FJ9-bound (red) ^15^N labelled (A) PDZ1 and (E) PDZ2 domains. (B, F) Δδ_avg_ of (B) PDZ1 or (F) PDZ2 in the presence of 10 molar equivalents of FJ9. The horizontal lines show the average Δδ_avg_ and the avg + stdev. Residues with the largest Δδ_avg_ are labelled. Syntenin 1 secondary structure, drawn from the PDB entry 1W9E(Grembecka et al., 2006b), is shown above the plot. (C, G) Chemical shift mapping of the FJ9 binding. The molecular surface of syntenin 1 (C) PDZ1 or (G) PDZ2 is colored by the Δδ_avg_ values (yellow: Δδ_avg_ > avg, orange: Δδ_avg_ > avg + stdev). Prolines and residues with unassigned backbone amide resonances are in grey. The docked FJ9 molecule is shown in sticks. (D, H) A detailed view of the docking solutions for FJ9 complexes with syntenin 1 (D) PDZ1 and (H) PDZ2 domains. Short intermolecular hydrogen bonds between FJ9 carboxyl group and the backbone amide protons of syntenin 1 are shown by dashed lines.

A low aqueous solubility of **FJ9** and its propensity to cause protein aggregation in NMR samples precluded a more detailed structural characterization of the syntenin-1/**FJ9** interaction. However, molecular modelling of **FJ9** binding to syntenin-1 PDZ1 and PDZ2 domains revealed physically plausible docking geometries, engaging the canonical peptide binding sites (Figure 4C-D,G-H). In both complexes, the **FJ9** molecule makes extensive intermolecular contacts, including numerous hydrophobic interactions and strong hydrogen bonds between the **FJ9** carboxyl group and backbone amide protons of PDZ1 I125 and G126 (Figure 4D) and PDZ2 V209 and G210 (Figure 4H). These are the key residues mediating the syntenin-1/**FJ9** interaction, as shown by our NMR analysis (Figure 4B-C,F-G) and further validated by molecular docking.

### Perturbation of the Tax-PDZ interactions and inhibition of Tax-1 functions using the small molecule FJ9

Next, we employed the YFP-based bimolecular fluorescence complementation (BiFC) assay, which provides direct visualization and quantification of protein interactions in living cells (Hu et al., 2002). We fused the two YFP-BiFC fragments to the *N-* and *C*-terminal ends of the Tax-1 and syntenin coding sequences, respectively. Pairs of fusion proteins were expressed in HEK293 cells, and fluorescent complementation was quantified by flow cytometry. As shown in Figures 5A-B, the interaction between Tax-1 and syntenin-2 exhibited the highest rate of complementation (33%), as compared to Tax-1 and syntenin-1 (14%). This finding is consistent with our co-immunoprecipitation and pulldown assays, showing stronger interactions with syntenin-2 (Figures 2F-G). To determine whether the small molecule **FJ9** could disrupt the Tax-1/syntenin interactions, HEK293 cells transfected with Tax-1/syntenin-1 or Tax-1/syntenin-2 pairs of YFP-BiFC fused constructs were treated with 300 μM of **FJ9** for 24h. **FJ9** treatment drastically diminished the rate of fluorescence complementation (33% to 4% for Tax-1/syntenin-2 and from 14% to 2.6% for Tax-1/syntenin-1) (Figures 5A-B). We therefore conclude that **FJ9** efficiently disrupts Tax/syntenin complex formation in cells.

**Figure 5.**
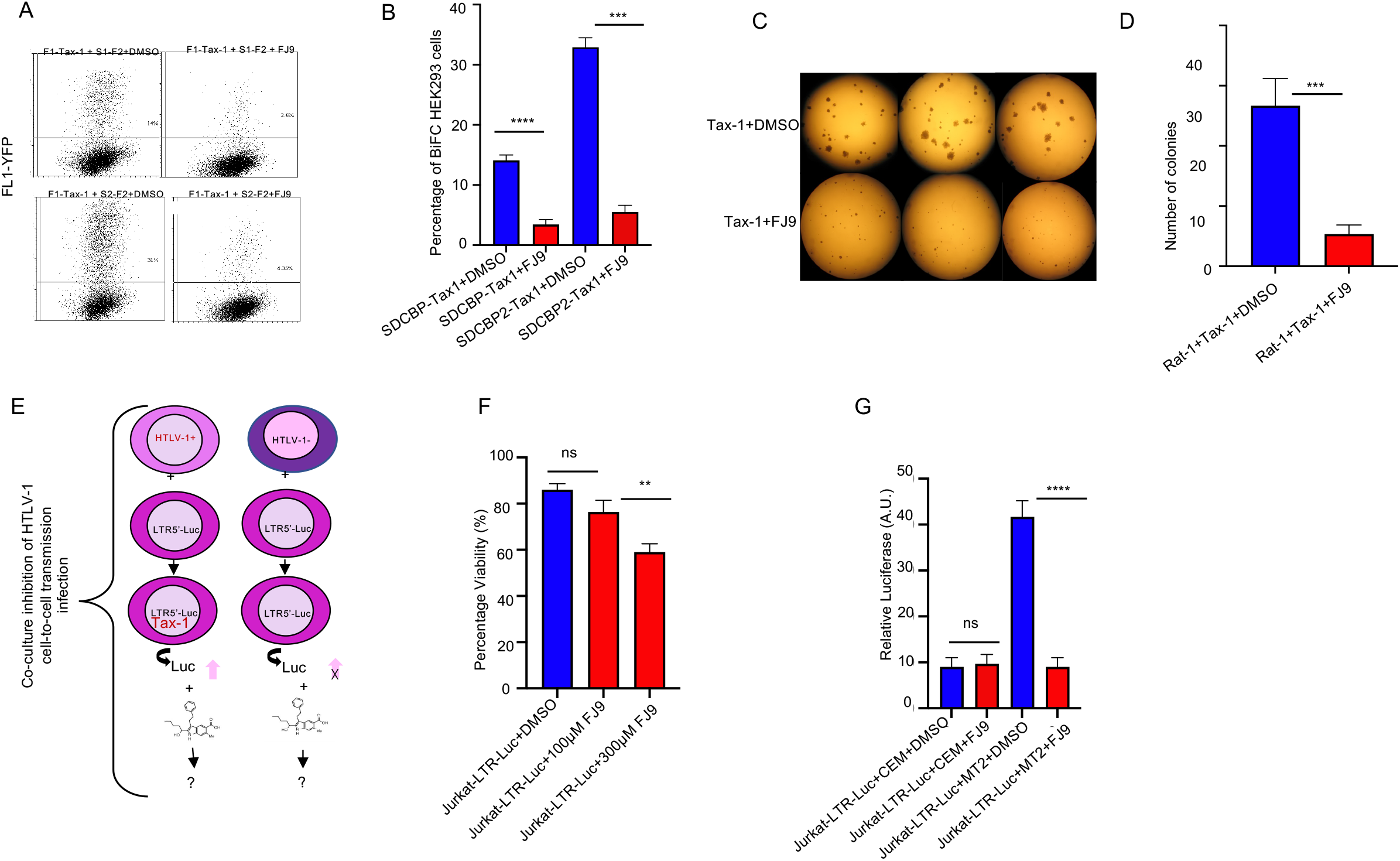
Perturbation of the Tax-PDZ interactions and inhibition of Tax-1 functions. (A) Positive HEK293 cells analyzed by FACS for fluorescence intensities in the presence (300 μM) or absence (DMSO) of the FJ9 small molecule. (B) Quantification of positive BiFC cells from (A). (C) Transformation of Rat-1 fibroblast cells following Tax-1 expression and treatment with 100 μM of FJ9 in a six-well plate. (D) Quantification of Tax-1-transformed colonies from (C), considering a minimum diameter cut-off of 0.25 mm. (E) Schematic representation of the co-culture infection and inhibition assay. (FF) Viability test, as measured using trypan blue, of Jurkat-LTR-Luc cells treated with DMSO, 100 μM or 300 μM of FJ9. (G) Quantification of cell-to-cell HTLV-1 transmission and inhibition by the FJ9 small molecule. Statistical analyses were done using unpaired t-tests, where **** indicate a p value <0.0001, *** a p value <0.001, and ** a p value <0.01.

The PBM of Tax-1 has been shown to increase Tax-1-mediated oncogenic capacity in cell culture (Higuchi et al., 2007). To examine whether **FJ9** could inhibit Tax-1 transformation activity, we infected a rat fibroblast cell line (Rat-1) with a VSV-G packaged lentiviral construct harbouring the HTLV-1 *tax* gene and the blasticidin resistant gene (CS-EF-IB-Tax-1). Cells were selected by blasticidin, and pools of resistant cells seeded for a colony formation in soft agar assay (CFSA) in the presence or absence of **FJ9** (100 μM). As shown in Figure 5C, HTLV-1 Tax transformed Rat-1 cells form multiple large colonies in CFSA, as previously described. However, in the presence of **FJ9**, Rat-1 cells formed much smaller colonies, and their number was reduced compared to the cells treated with the vehicle. Thus, we conclude that **FJ9** inhibits Tax-transformation activity in the Rat-1 model (Figures 5C-D).

To probe the role of the Tax/PDZ interaction in cell-to-cell HTLV-1 transmission, we performed a co-culture experiment using Jurkat T-cell line reporter expressing luciferase under the control of the 5’LTR HTLV-1 promoter, and a HTLV-1 producing cell line (MT2) or as negative control, a human T cell leukemia cell line (CEM) (Figure 5E). The reporter exhibits basal levels of luciferase expression, which increase following co-culture with MT2 cells and transactivation of the 5’LTR by Tax. We show that at 100 μM, a dose which does not induce significant cell death compared to the vehicle (Figure 5F), **FJ9** was able to inhibit HTLV-1 transmission, reducing Tax transactivation to the basal level (Figure 5G).

### Perturbation of Tax-1 localization in the Golgi apparatus and virological synapses by the small molecule FJ9

The cell-to-cell transmission of HTLV-1 depends on the polarization of intracellular organelles, including the Golgi apparatus, towards virological synapses (Barnard et al., 2005; Igakura et al., 2003; Nejmeddine et al., 2009). To better understand the cellular effects of **FJ9** during HTLV-1 transmission, we co-immunostained MT2 cells with antibodies targeting a cis-Golgi marker (GM130) and the HTLV-1 Tax protein (Figure 6A). Using 3D reconstructions from z stack series of confocal microscopy images, we showed that **FJ9** treatment affects Golgi apparatus organization by increasing its intracellular volume (Figure 6B, GM130 staining, p-value<0,0001). In addition, immunostaining of Tax-1 showed that **FJ9** increases the number of Tax-1 cytoplasmic foci, characterized by heterogenous volume ranging from 20 to 150 μm^3^ (Figure 6C, GM130 p-value<0,01). Moreover, the mean fluorescence intensity associated with Tax-1 expression significantly increased in GM130 positive clusters following **FJ9** treatment, suggesting that the intracellular distribution of Tax-1 is affected. We also tested the effect of **FJ9** on virological synapse formation. To this end, MT2 cells were cultivated with non-infected Jurkat cells at a 1:1 ratio for 24h in presence or absence of **FJ9**. Additionally, we analyzed the cells using confocal microscopy and showed that MT2 cells are able to perform cell-cell interactions with Jurkat cells. Virological synapses are identified when MT2/Jurkat interaction is accompanied by segregation of CD11a and Tax-1 expressed by MT2 at the contact area with a non-infected Jurkat cell (Figures 6E and 6F). Our data show that the frequency of synapses are increased by two-fold when MT2 are co-cultured with Jurkat cells in the presence of **FJ9** (average 27% to 61% of conjugates, p-value<0,05) (Figure 6G). Taken together, our results demonstrate that **FJ9** impairs the intracellular localization of Tax-1 but maintains polarized vesicular bodies at synapses.

**Figure 6.**
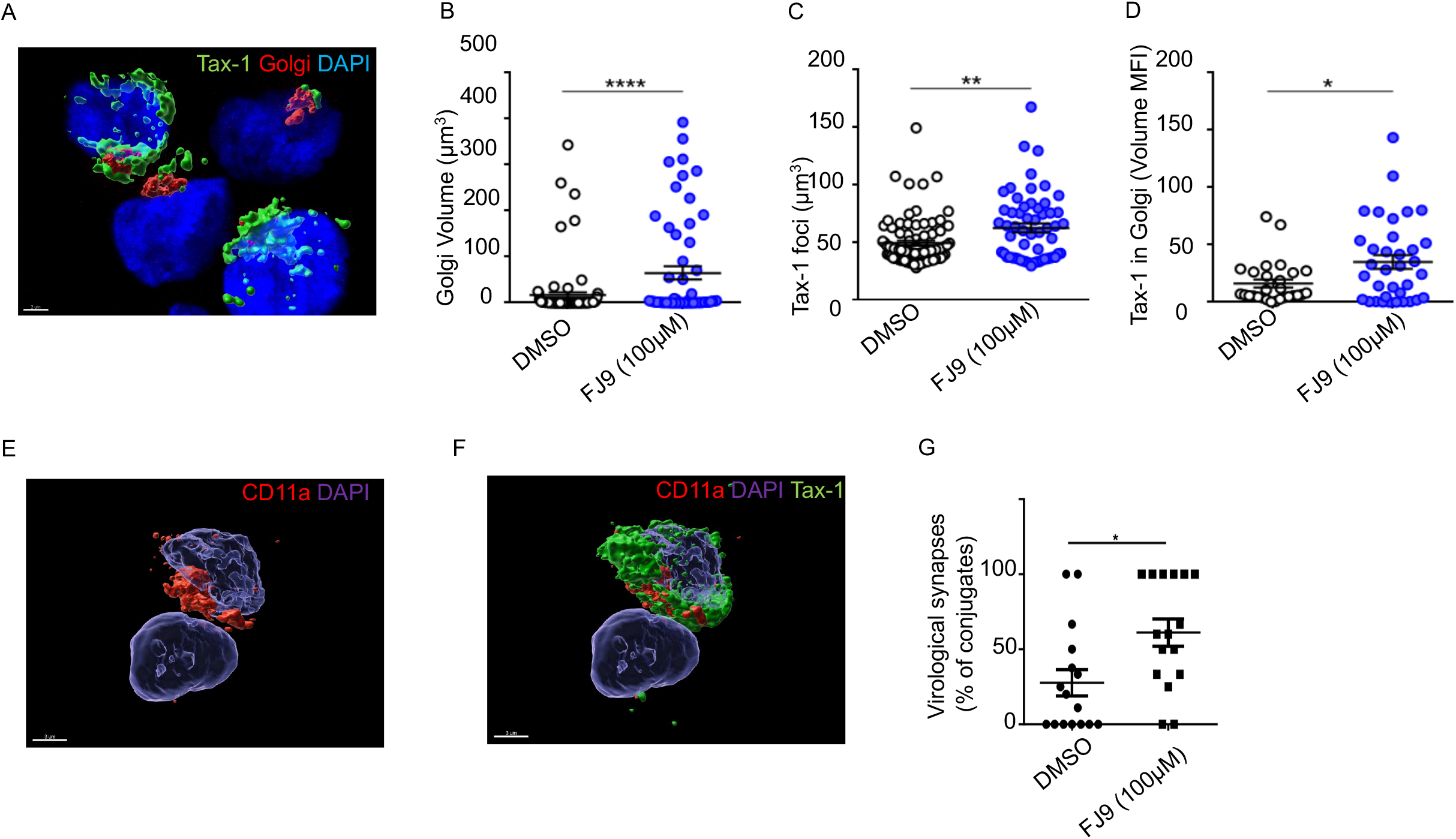
Perturbation of Tax-1 localization in the Golgi apparatus and virological synapses by FJ9 molecule. (A) Co-immunostaining of MT2 cells with cis-Golgi marker (GM130) antibodies and Tax-1 protein. (B) Golgi apparatus intracellular volume. (C,D) immunostaining of Tax-1, p-value <0.01. (E, F) virological synapses identified when MT2/Jurkat interaction is accompanied by segregation of CD11a and Tax-1 expressed by MT2 at the contact area with non-infected Jurkat cell. (G) Frequency of synapses in the presence of FJ9, p-value <0.05

## DISCUSSION

Despite the discovery of viruses in the early 1900s, and the demonstration of their pathogenicity in humans in early 1950s, the arsenal of antiviral drugs remains dangerously small, with only ~90 molecules available today (“Principles of Virology, Volume 2,” n.d.). Those are biased towards viral enzymes, including polymerases, proteases, and integrases. Furthermore, 95% of approved drugs target only 6 types of viruses (HIV-1, HCV, HBV, HSV, HCMV, and Influenza viruses), with HIV-1 covering the majority of available drugs (De Clercq and Li, 2016). High-throughput mapping of host-virus PPIs and massive parallel genomic sequencing have accelerated the pace of key host target discovery, which are shared by different categories of pathogenic viruses, including RNA and DNA, or acute and persistent viruses (Rozenblatt-Rosen et al., 2012; Tang et al., 2013). Identified host factors provide novel opportunities to prioritize antiviral drugs based on common targets (Olivet et al., 2022). In this work, our goal was to demonstrate that one type of such host determinants, the PDZ-domains, could provide insights into how to discover small molecule inhibitors of viral-host protein-protein interactions.

By testing 248 out of 271 PDZ domains identified in 151 human proteins, our results establish the first comprehensive Tax-1 /PDZ interactome, highlighting a set of 54 PDZ-domain-containing proteins. Comparative analyses with similar unbiased mapping for other viral proteins are now possible. For example, hDLG1 and SCRIB have been shown to interact with HPV E6, Ad9 E4orf1, IAV NS1, HTLV-1 Env, HIV-1 Env and HCV core proteins, but with different biological outcomes, including perturbations of cell polarity, signaling, or apoptosis (Thomas and Banks, 2018). Interestingly, the conserved *C*-terminal PBM motifs of these viral proteins highly correlate with viral pathogenicity, independently of their substantial differences in genome content (RNA and DNA viruses) or mode of infection (acute and persistent families of viruses). Other examples include syntenin-1 and PTPN13, both able to interact with distant viral proteins such as the HTLV Tax-1 (this study), but also with the coronaviral E, 3A and N proteins (Caillet-Saguy et al., 2021; Caillet-Saguy and Wolff, 2021; Jimenez-Guardeño et al., 2014). These pan-viral interactors may thus constitute ideal drug targets for the discovery of broad-spectrum antiviral inhibitors. Consistent with the above findings, we previously highlighted the importance of syntenin-1 as a potential recruiter of proteins into extracellular vesicles (EV) (Lim et al., 2021). Here, we further demonstrate that, through its PDZ domains, syntenin-1 recruits Tax-1 into EV (Figure 2B), which provides a mechanistic explanation for the presence of the Tax-1 protein in EVs isolated from HTLV-1-infected cell lines, HAM/TSP or ATL patient samples (Barclay et al., 2017; Jaworski et al., 2014; Otaguiri et al., 2018). A compelling hypothesis is that blocking syntenin-1/viral proteins interactions could elicit a two-pronged inhibitory activity: (i) at the viral budding step of enveloped virus replication, where syntenin-1 could connect viral particles to the ESCRT pathway (Göttlinger et al., 1991; Howard et al., 2001; Votteler and Sundquist, 2013) and (ii) in cell-to-cell viral transmission and inflammatory immune response induction by exosomes containing syntenin-1/viral proteins complexes (Otaguiri et al., 2018). Interestingly, our study shows a correlation between inhibition of syntenin-1 /Tax-1 interaction and redistribution of Tax-1 in the Golgi apparatus (Figures 5A-B and 6A-D). In support of the findings described above, a previously described small-molecule inhibitor of the syntenin-exosomal pathway was also shown to potentially disrupt tumor exosomal communication in breast carcinoma cells (Leblanc et al., 2020), suggesting that targeting the presence of viral or cellular oncogenes in EV could affect their transport between cells, and provide an alternative inhibition strategy.

The structural analysis performed in this work shows that the peptide corresponding to the *C*-terminal segment of Tax-1 protein binds PDZ1 and PDZ2 domains of syntenin-1. This Tax-1 10-mer engages the canonical peptide binding sites of both domains (Figure 3), which is consistent with the binding mode observed in other syntenin-1/peptide complexes (Grembecka et al., 2006; Latysheva et al., 2006). The binding effects in the PDZ1+2 tandem are more pronounced for the PDZ2 domain, as also seen in the earlier NMR study of syntenin-1 interactions with syndecan, neurexin, and ephrin B-derived hexapeptides (Grembecka et al., 2006).

Given that PDZ domains recognize small linear motifs on cognate protein targets, the use of short peptides to mimic protein-PDZ interactions is a valid and ubiquitous experimental strategy to unravel relevant physiological interactions for fundamental research purposes (Manjunath et al., 2018), but it has its limitations. In particular, biophysical assays performed in dilute solutions of isolated syntenin-1 PDZ domains and short peptide ligands report very weak binding with K_D_ values in the mM range (Figure 3) (Grembecka et al., 2006; Latysheva et al., 2006), which is clearly insufficient to sustain a physiological interaction. However, in the context of the full-length proteins in the cellular millieu, the Tax-1/syntenin-1 binding is expected to be much stronger (as confirmed in this work by BiFC and pull-down assays). In particular, a cooperative interaction of the native Tax-1 homodimer with syntenin-1 PDZ tandems, possible contributions of other protein domains to the Tax-1/syntenin-1 binding, and crowding and confinement effects, known to enhance biomolecular association *in vivo* (Bernadó et al., 2004; Phillip and Schreiber, 2013), could account for the discrepancy in the observed binding strengths.

As demonstrated in this work, the described small molecule **FJ9** clearly targets the peptide binding sites of both PDZ1 and PDZ2 domains of syntenin-1 (Figure 4), which explains its inhibitory effect on Tax-1 association. The absolute values of the NMR chemical shift perturbations and the extent of the **FJ9** binding effects were smaller than those of the Tax-1 10-mer (cf. Figures 3 and 4), most likely reflecting a set of less tight contacts made by the small molecule ligand. These findings are in line with an NMR study of a Dishevelled protein PDZ domain, which also reported a large decrease of **FJ9** binding effects as compared to those of the cognate peptide (Fujii et al., 2007). Overall, our molecular docking models of **FJ9** complexed with the syntenin-1 PDZ domains agree with the NMR chemical shift perturbation data and capture particularly well the largest binding effects (backbone amides of I125, G126, V209, and G210; Figure 4). Importantly, the presented molecular models could be valuable starting points for the future structure-based design of more potent inhibitors of the Tax-1/syntenin-1 interaction. Along the same lines, further elaboration or chemical modification of small molecule inhibitor hit or lead compounds would be advantageous over peptide-based inhibitors, as the former will generally be characteristic of a higher *in cellulo* access, and hence eventual therapeutic outcomes.

Direct antivirals against viral enzymes efficiently attenuate viral replication. However, resistant mutations arise rapidly, suggesting that inhibitors targeting virus-host interactions should synergize the antiviral potential and minimize the occurrence of drug resistant mutants. Developing this class of inhibitors come with additional challenges including (i) the identification of “hot spots” i.e. amino acid residues contributing most to the binding energy of two protein partners, and (ii) potential side effects that could arise from blocking other functions of the targeted host proteins. These might be overcome by (i) the early determination of three-dimensional structures of the PPI targets, (ii) the design and optimization of low-molecular weight and highly selective peptidomimetic compounds based on interacting domains and motifs, and (iii) by the early-stage testing of molecules in relevant cell lines and animal models. The above steps might allow to lower the probability of the appearance of resistance mutations, and would potentially still allow natural ligands to interact with the endogenous target protein. Complete inhibition of a specific host target function required for viral spreading is also possible and would mirror gene knock out of host dependency factors, as exemplified in studies using the CRISPR technology (Kirby et al., 2021; Li et al., 2020; Watanabe et al., 2014).

In conclusion, just as the identification of the main immunogenic proteins of a pathogenic virus allows rapid development of candidate vaccines, systematic interactome PPI mapping is of utmost importance to delineate druggable virus-host PPIs. We have demonstrated a clear correlation between PPI inhibition and impairment of HTLV-1 Tax functions, including its transformation ability and HTLV-1 cell-to-cell transmission through the exosomal pathway. The combination of our experimental strategies should allow the development of more potent anti-Tax-1 compounds for future pre-clinical and clinical studies of drug candidates in view of treating ATL and other HTLV-1-induced diseases.

## AVAILABILITY

All data and materials used in the analyses are described in this manuscript. Plasmids and vectors are available via materials transfer agreements (MTAs).

## SUPPLEMENTARY DATA

Supplementary Data are available at JER online.

## ACKNOWLEDGEMENT

We thank Dr. Fujii Naoaki (Purdue University) for providing the **FJ9** compound. Computational resources have been provided by the Consortium des Équipements de Calcul Intensif (CÉCI), funded by the Fonds de la Recherche Scientifique (FRS-FNRS, Belgium) under Grant No. 2.5020.11 and by the Walloon Region. We thank the Cell Imaging and Viral Vectors core facilities of the University of Liege for their services.

---

**Figure S1.**
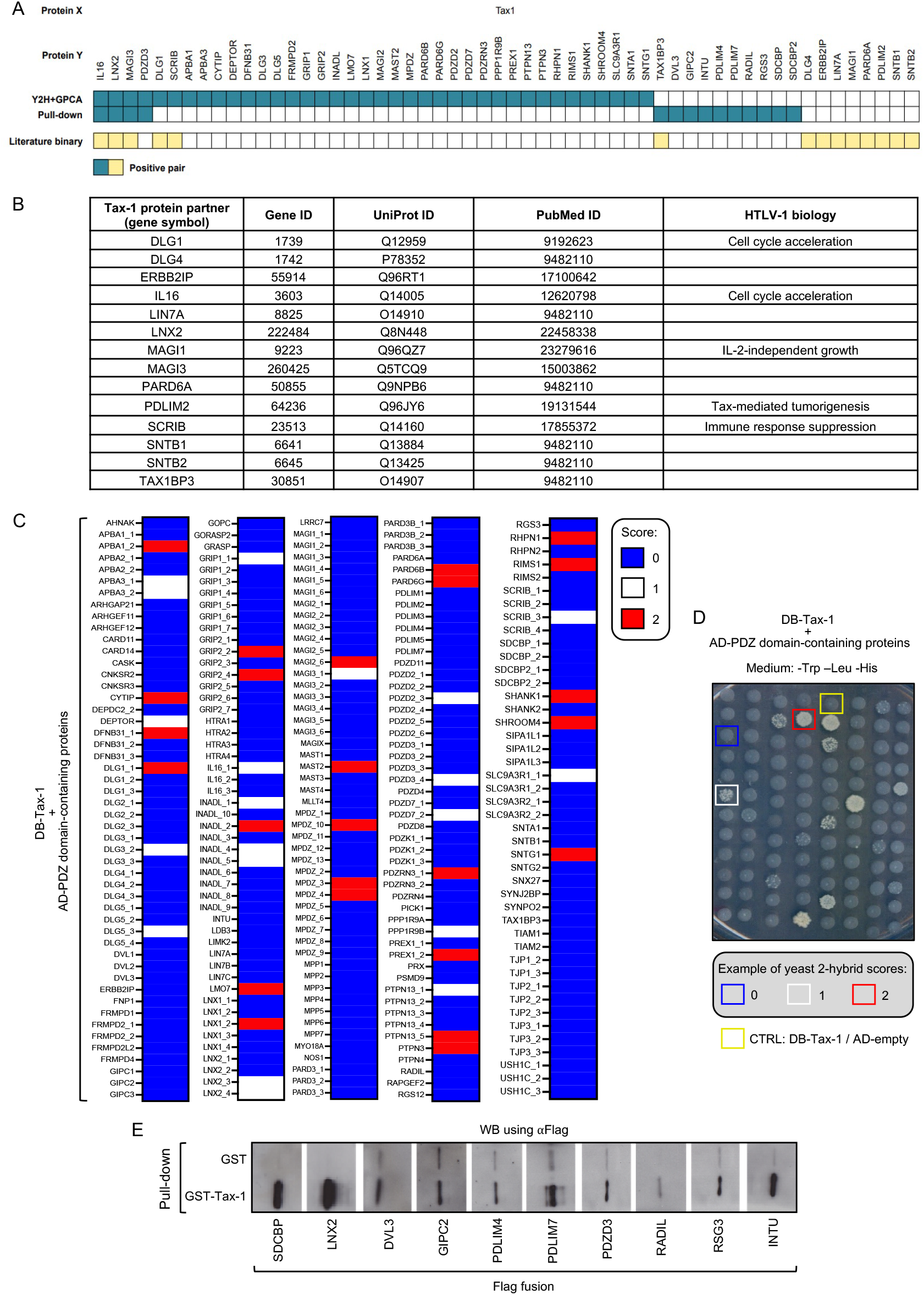
A comprehensive analysis of the Tax-1 interactome with human PDZ-containing proteins. (A) PPIs between Tax-1 and human PDZ domain-containing proteins identified by yeast 2-hybrid (Y2H) and validated by GPCA or using a pull-down assay (blue); and known PPIs from the literature (yellow). (B) Literature reported interactions between Tax-1 and human PDZ domain-containing proteins. (C) Heat maps of the Y2H assay by testing a library of 244 individualized PDZ domain-containing proteins fused to AD against DB-Tax-1. A growth score of “0”, “1” or “2” indicates a null, weak or strong interaction, respectively. (D) Representative Y2H assay plate lacking tryptophan (Trp), leucine (Leu), and histidine (His) where a growth score of “0”, “1” or “2” indicates a null, weak or strong interaction between DB-Tax-1 and PDZ-domain-containing proteins fused to AD, respectively. (E) Representative GST-pull-down analyses of proteins encoded by Flag-fused open reading frames (ORFs) available in the human ORFeome collection.

